# Purinergic Preconditioning Induces Epigenomic and Transcriptomic Changes Resembling Epilepsy-associated Microglial States

**DOI:** 10.1101/2023.06.21.545837

**Authors:** Ricardo Martins-Ferreira, Josep Calafell-Segura, João Chaves, Laura Ciudad, António Martins da Silva, Paulo Pinho e Costa, Bárbara Leal, Esteban Ballestar

## Abstract

Microglia, as the main immune effector cells in the central nervous system (CNS), play a crucial role in a diverse range of neuropathological conditions through their exacerbated activation. Microglial inflammatory responses can be influenced by prior exposures to noxious stimuli, such as increased levels of extracellular adenosine and ATP. These conditions are characteristic of brain insults like epileptic seizures and could potentially shape subsequent responses through epigenetic regulation. In this study, we investigated DNA methylation and expression changes in microglia-like cells differentiated from monocytes following ATP-mediated preconditioning. First, during differentiation, we demonstrate that microglia-like cells acquire standard microglial features, including whole transcriptomes and methylomes like *in vivo* profiles. We show a predominant DNA demethylation in genomic sequences enriched in binding motifs of microglia lineage transcription factors like PU.1, consistent with the relevance of this factor in *in vivo* microglia. TLR-mediated activation, after a first exposure to ATP, promotes exacerbated pro-inflammatory activation compared to cells not pre-exposed to ATP. These changes are accompanied by DNA methylation and transcriptional reprogramming associated with the acquisition of trained immunity and altered immune-related functions such as with antigen presentation, phagocytosis and cytokine signaling. Finally, the reprogramming associated with ATP-mediated preconditioning leads to profiles found in microglial subsets linked to epilepsy. Purine-driven microglia immune preconditioning drives epigenetic and transcriptional changes that could contribute to altered functions of microglia during seizure development and progression, particularly associated with neuroinflammation.

## INTRODUCTION

Microglia, as the brain’s resident macrophage population, represent the first line of immune defence within the central nervous system (CNS). Despite their low abundance in relation to other cell types (around 10%) (1,2), consistent evidence has demonstrated an essential role of microglia in maintaining overall brain function (3–5). Furthermore, mutations of microglia-specific genes have been associated with different neurodegenerative diseases (6,7). The involvement of microglia in neuropathology is often attributed to its aberrant activation. Recently, exacerbated and uncontrolled pro-inflammatory activation has been attributed to the existence of innate immune memory (8). Similar to what has been demonstrated in other myeloid cells, like monocytes and macrophages, microglia are prone to respond differently to secondary stimuli when pre-exposed to a first stimulus, in a process tightly regulated by epigenetic reprogramming (9–12).

The study of purinergic signalling may represent a promising pathway to better understand the role of microglial conditioning in epileptogenesis. Adenosine and ATP have been proposed to be essential mediators of epilepsy (13). However, there is no consensus on the precise underlying mechanisms that are involved. Generally, adenosine is primarily considered under the light of its anticonvulsant effects through activation of adenosine A1 receptors (14), and may also exert excitatory and neurodegenerative effects associated with adenosine A2A receptor activation (15,16). Like adenosine, ATP is a ubiquitous endogenous molecule with multiple implications on the CNS through modulation of cell survival, proliferation and differentiation, axonal growth and maturation, excitability and glial activation (17). The impact of ATP in epileptogenesis has mainly been studied in the context of the ionotropic P2X7 receptor (P2RX7), the activation of which is predominantly associated with microglia activation and pro-inflammation. P2RX7 is generally increased in epilepsy with anticonvulsive effects attributed to its antagonistic treatment (13). Purine metabolism is widely pleiotropic, with the respective receptors being expressed throughout all the major cell types in the CNS, and therefore exerting potentially contrary effects depending on the cell type, the receptor, the brain region, the used model of epilepsy and genetic heterogeneity. Understanding in depth the intricacies of the impact of purinergic imbalance in epileptogenesis may represent an overwhelming task. What appears to be more widely accepted is that both adenosine and ATP extracellular levels increase significantly upon brain insult, including after the occurrence of seizures. It is also important to take into consideration that ATP has a short half-life and is quicky converted into adenosine by ectonucleotidases. Therefore, the increased release of ATP also contributes to elevated levels of adenosine (18–21).

Human microglia research has been significantly hampered by the low replicability of *in vivo* traits in cell culture models. The use of foetal or adult brain biopsies for human primary microglia cultures is limited by ethical and logistic reasons, together with relatively low yields of isolation (22). In addition, replicating the homeostatic microglia phenotype is particularly challenging due to the substantial transcriptomic and epigenetic changes that cells undergo in culture (23), and may be influenced by post-mortem conditions (24). To overcome these limitations in experimental scalability of human microglia cultures, alternative protocols have been developed, including the differentiation of peripheral blood monocytes into microglia-like cells. In this study, we aimed to investigate how ATP-driven preconditioning can influence posterior inflammatory activation in microglia by using monocyte-derived microglia-like cells. In our study, we determined the DNA methylation and transcriptional changes associated with inflammatory preconditioning, the associated features and its potential relationship with the phenotype of microglia in epileptogenesis.

## RESULTS

### Characterization of monocyte-derived microglia-like cells

We first differentiated microglia-like cells from peripheral blood monocytes using an adapted version of a previously described procedure (25). Monocytes were differentiated in serum-free culture medium supplemented with M-CSF, GM-CSF, β-NGF, IL-34 and CCL2. Moreover, we added TGF-β2 and cholesterol solution, which have been shown to increase viability of mice microglia *in vitro* (26) (Figure 1A). In parallel, we differentiated monocytes to pro-inflammatory macrophages as a reference to validate the homeostatic nature of microglia-like cells. Microglia-like cells acquired a stable phenotype at seven days. After three days, we observed that cells acquired an elongated and ramified morphology resembling homeostatic microglia *in vivo* (Figure 1B). Gene expression evaluation by real-time (RT)-PCR demonstrated upregulation of canonical homeostatic microglia markers *CX3CR1* and *P2RY12* in microglia-like cells in comparison to monocytes (Figure 1C). We also performed flow cytometry evaluation of a panel of markers consistently used to describe and distinguish microglia-like from monocytes and macrophages. Our gating strategy aimed to delimitate the CD11b+ macrophage lineage population in the microglia-like and monocyte-derived macrophages (Supplementary Figure 1A). We observed a decrease in CD14 and CCR2 in relation to monocytes and macrophages, as previously described (27); an increase in CD68, a standard microglia marker in brain tissue besides also being expressed by other macrophages, in relation to monocytes; and a decrease in CD45 in relation to macrophages (Figure 1D and Supplementary Figure 1B). It is important to note that the dual CD11b/CD45 signal is the most predominantly used strategy to distinguish homeostatic microglia (CD11b^+^/CD45^low^) from macrophages (CD11b^+^/CD45^high^), and this model replicated these conditions.

**Figure 1.**
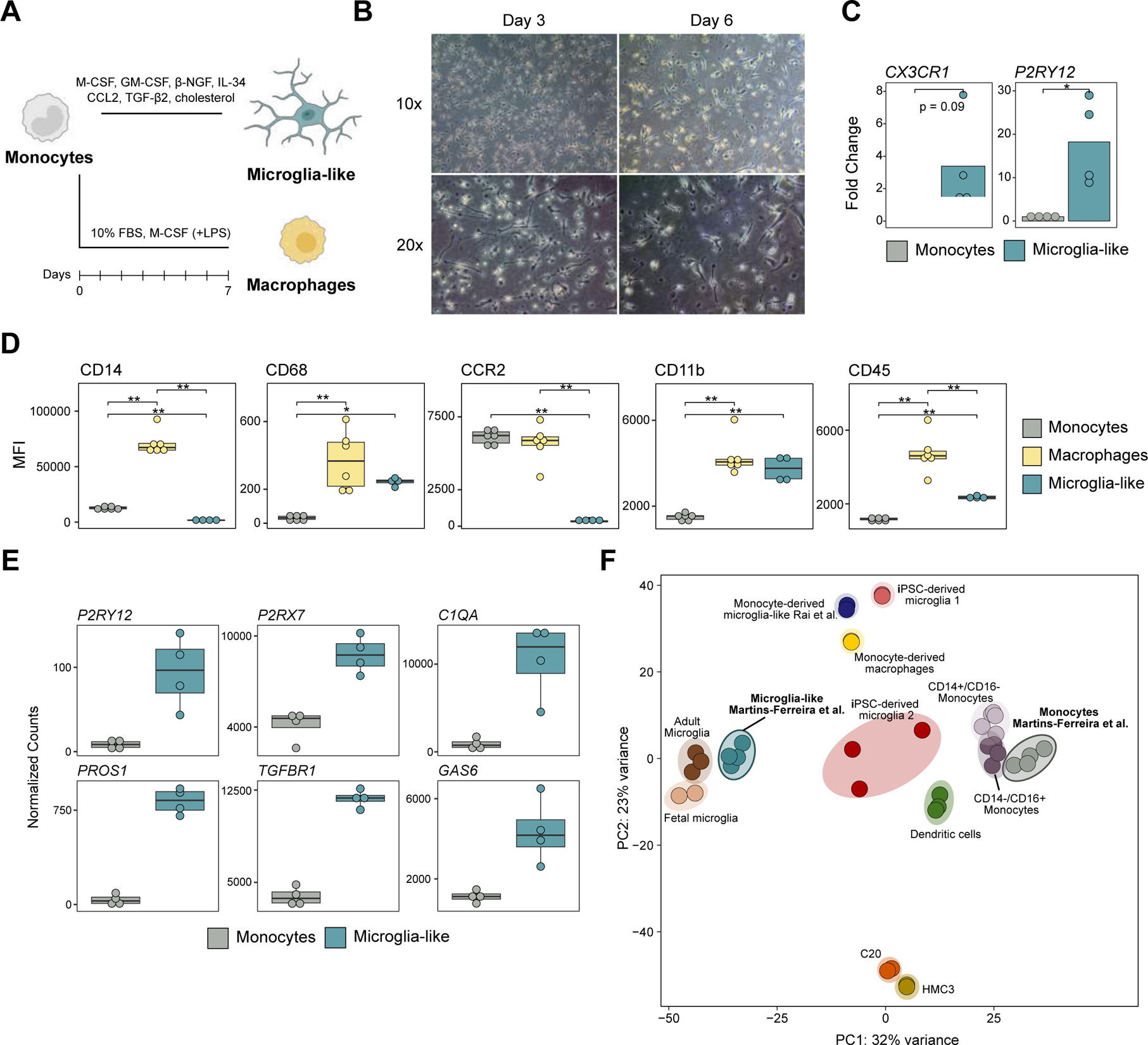
**(A)** Schematic representation of the *in vitro* differentiation protocol from freshly isolated monocytes to monocyte-derived microglia-like cells and monocyte-derived macrophages. For microglia-like differentiation, monocytes were plated in serum-free conditions with medium supplemented with M-CSF, GM-CSF, β-NGF, IL-34, CCL2, TGF-β2 and cholesterol. Macrophages were obtained by culturing monocytes in medium supplemented with 10% FBS and M-CSF; plus, LPS treatment one day prior to collection. **(B)** Representative optical microscopy images of microglia-like differentiation at days three and six of culture, with 10x and 20x magnification. **(C)** Barplot representation of the gene expression of *CX3CR1* and *P2RY12* in microglia-like cells in comparison to monocytes (*p<0.05). **(D)** Boxplot representation the median fluorescence intensity (MFI) values obtained by flow cytometry for CD14, CD68, CCR2, CD45 and CD11b in monocytes, macrophages and microglia-like cells *p<0.05, **p<0.01). **(E)** Boxplot representation of RNA-seq normalized counts for *P2RY12*, *P2RX7*, *C1QA*, *PROS1, TGFBR1* and *GAS6* in monocytes and microglia-like cells. All four genes were significantly upregulated in microglia-like cells in the regression model. **(F)** Principal Component Analysis (PCA), using Variance Stabilizing Transformation (VST) values considering all transcriptome of the RNA-seq data generated in this study for monocytes and microglia-like, together with public data of CD14+/CD16- and CD14-/CD16+ monocytes, dendritic cells, monocyte-derived macrophages, monocyte-derived microglia-like, iPSC-derived microglia, and primary adult and fetal microglia from Rai *et al.* (2020).

To further assess the acquisition of the resting microglia phenotype at the transcriptomic level, we performed RNA-seq analysis of monocytes and microglia-like cells. Differentiation of monocyte to microglia-like cells resulted in the upregulation of 6500 genes and downregulation of 5847 genes (FDR<0.05) (Suppl. Table 1). For instance, we observed upregulation of *P2RY12*, *P2RX7*, *C1QA*, *PROS1, TGFBR1* and *GAS6* (Figure 1E, Supplementary Figure 2A and Supplementary Table 1), microglia markers previously shown increased in microglia-like cells (25). We integrated our RNA-seq data with public datasets corresponding to microglia and other myeloid populations (Figure 1F). These were described by Rai *et al.* (2020) (28), and included CD14+/CD16-(classical) and CD14-/CD16+ (non-classical) monocytes, conventional dendritic cells, two sets of induced pluripotent stem cell (iPSC)-derived microglia, monocyte-derived macrophages, the monocyte-derived microglia-like cells generated in that study, microglia cell lines (C20 and HMC3) and primary human adult and foetal microglia. We observed that our monocytes clustered together with those from the other datasets, and that the microglia-like cells generated with our method clustered closer to adult and foetal microglia. These results demonstrate the validity of microglia-like cells to replicate microglial transcriptomic features.

### DNA methylation changes during differentiation associate with the acquisition of microglial features

We next determined the DNA methylation changes associated with the differentiation to microglia-like cells. In comparison to monocytes, homeostatic microglia-like cells had 2879 differentially methylated positions (DMPs) (FDR < 0.05 and absolute mean Beta difference > 0.2) (Supplementary Table 2). Most DMPs (2799) corresponded to CpG sites that are hypomethylated in microglia-like cells in comparison with monocytes (Figure 2A). Gene ontology (GO) analysis of hypomethylated DMPs revealed enrichment of terms associated with inflammation and leukocyte differentiation (Figure 2B). Moreover, transcription factor (TF) binding motif enrichment analysis yielded multiple TFs associated with the myeloid lineage (Figure 2C). Within those, we were able to identify several TFs previously associated with microglia-specific epigenetic modelling in human brain tissue [23,29], a list headed by PU.1 (Figure 2D). To further validate the microglial nature of our microglia-like cells, we integrated our DNA methylation profiles with public data from microglia isolated from 56 human brain tissue samples (GSE191200). We observed that microglia-like cells cluster closer to primary microglia samples than to monocytes (Figure 2E). Integration of DNA methylation and RNA-seq showed that the genes associated with hypomethylated DMPs were upregulated in microglia-like cells vs monocytes (Figure 2F). Selected examples of genes displaying both hypomethylation near the TSS and upregulation during microglia-like differentiation include *CD68* and *CSF1R,* which are implicated in differentiation within the macrophage lineage and are essential for microglia maintenance and survival *in vivo* (30) (Figure 2G).

**Figure 2.**
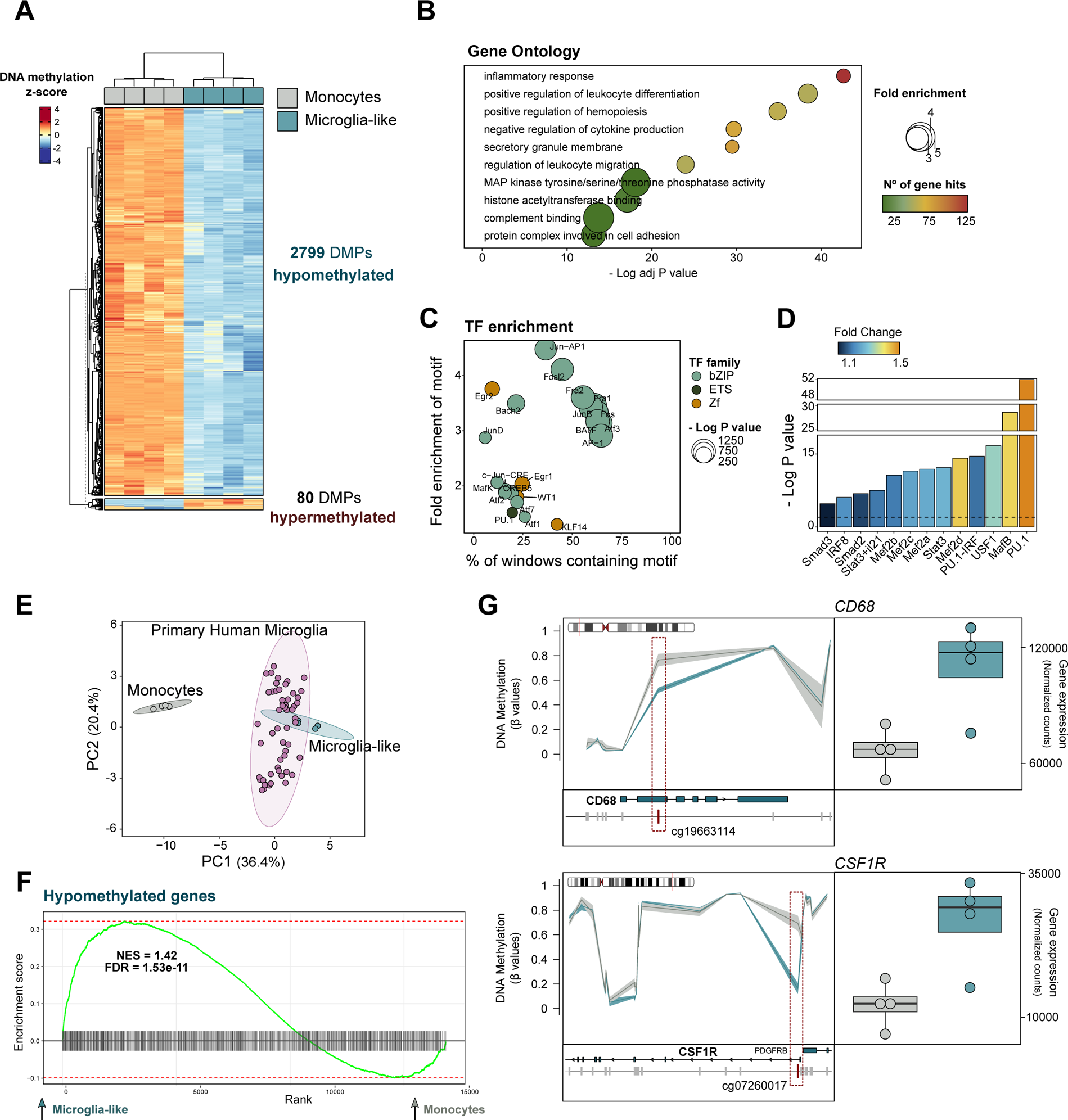
**(A)** Heatmap representation of the DNA methylation of the differentially methylated positions (DMPs) obtained for the microglia-like vs monocytes comparison. DNA methylation is represented as the z-score of the beta values. The significance cutoff was of FDR < 0.05 and difference in mean beta values > 0.2. **(B)** Selected list of significantly enriched gene ontology (GO) terms for the hypomethylated DMPs in microglia-like. The significance of enrichment is represented by the negative of the logarithm of the adjusted p value, the enrichment fold change and the number of gene hits. **(C)** Motif enrichment of the most significant of the most significant transcription factors (TFs) (p value < 1E-21). The TFs are annotated by family and the enrichment is represented by the negative of the logarithm of the p value. **(D)** Barplot showing the TF motif enrichment of a selected group of TFs previously show to be associated with microglia-specific chromatin accessibility. All show a p value lower than 0.01. **(E)** Principal Component Analysis (PCA) considering the beta values corresponding to all pairwise DMPs between monocytes, microglia-like cells and primary microglia from post-mortem brain (GSE191200). **(F)** Gene Set Enrichment Analysis (PCA) of the list of genes associated with hypomethylated DMPs in the differential expression comparison between microglia-like and monocytes. Genes associated with hypomethylated DMPs are upregulated in microglia-like cells. The enrichment is represented by the positive Normalized Enrichment Score (NES) and the statistical significance by the FDR. **(G)** Graphical representation of the beta values of the CpGs located nearby the *CD68* and *CSF1R* for monocytes and microglia-like cells (left panels). The genes and the individual probes are represented in relation to the annotated genes in the UCSC Ref Seq. The CpGs highlight in red demonstrate a statistically significant hypomethylation in microglia-like vs monocytes. Boxplot representation of RNA-seq normalized counts for *CD68* and *CSF1R* in monocytes and microglia-like cells (right panels). Both genes were significantly upregulated in microglia-like cells in the regression model.

### ATP-driven immune preconditioning results in epigenetic and transcriptional remodeling in microglia-like cells

To interrogate whether an initial ATP stimulus influences microglia responses to secondary inflammatory activation, we stimulated microglia-like cells with ATP and then subjected them to TLR4-mediated activation with lipopolysaccharide (LPS) (Figure 3A). To this end, after 6-day microglia differentiation, we treated a sample with ATP one day before harvesting (preconditioned), whereas in parallel we left a sample without ATP treatment (non-preconditioned). In addition, we performed microglia activation using LPS in both microglia samples, preconditioned and non-preconditioned with ATP, and collected the samples after two days. The experiments were conducted using four biological replicates. Firstly, we observed upregulation of pro-inflammatory genes *IL1B* and *IL6* in activated preconditioned microglia in comparison with activated non-preconditioned (Figure 3B). IL-1β protein levels were also increased in the supernatant of activated preconditioned microglia (Figure 3C). The exacerbated pro-inflammatory profile of preconditioned microglia-like cells was corroborated by flow cytometry. They showed higher levels of CD14, CD68, CD45 and HLA-DR in comparison with activated non-preconditioned microglia (Figure 3D and Supplementary Figure 3A).

**Figure 3.**
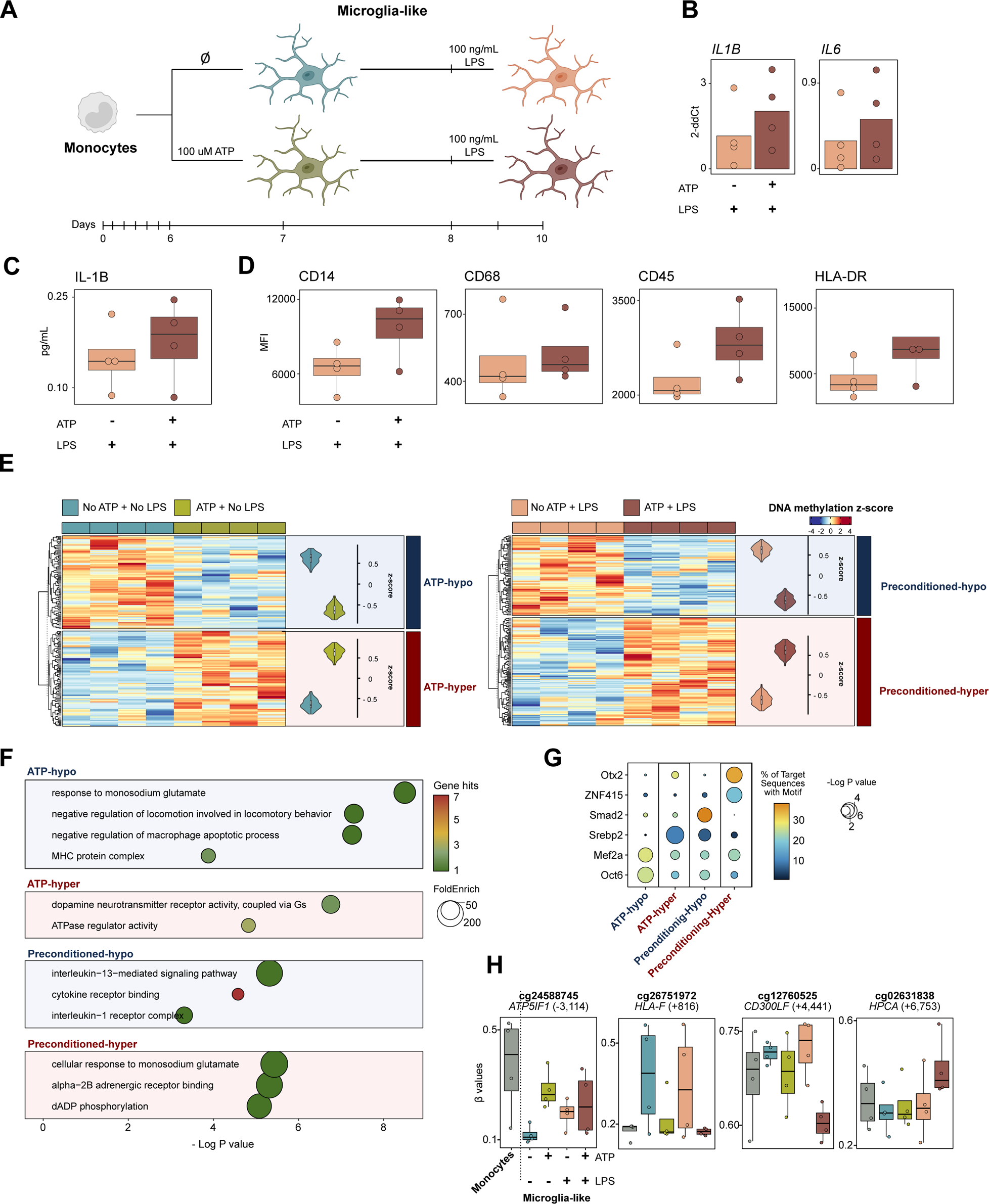
**(A)** Schematic representation of the preconditioning and inflammatory stimulation strategy in microglia-like cells. The first set of conditions were collected at day seven of culture and were either preconditioned or non-preconditioned with 100 µM ATP for 24 hours. The preconditioned and non-preconditioned microglia-like were stimulated with 100 ng/mL LPS two days before collection and collected at day ten of culture. **(B)** Barplot representation of the gene expression of *IL1B* and *IL6* in microglia-like activated preconditioned vs non-preconditioned (*p<0.05) obtained by RT-PCR. **(C)** Boxplot representation of protein levels of IL-1B in the supernatant of microglia-like activated preconditioned vs non-preconditioned. **(D)** Boxplot representation the median fluorescence intensity (MFI) values obtained by flow cytometry for CD14, CD68, CD45 and HLA-DR in microglia-like activated preconditioned vs non-preconditioned. **(E)** Heatmap representation of the DNA methylation of the differentially methylated positions (DMPs) obtained for the preconditioned vs non-preconditioned in activated and non-activated microglia like cells. DNA methylation is represented as the z-score of the beta values. The significance cutoff was of p value < 0.05 and absolute difference in mean beta values > 0.1. The violin plots represent the mean z-score of each methylation site in each group. **(F)** Selected list of significantly enriched gene ontology (GO) terms for each of the lists of DMPs (*ATP-hypo*, hypomethylated in non-activated preconditioned vs non-preconditioned; *ATP-hyper*, hypermethylated in non-activated preconditioned vs non-preconditioned; *Preconditioned-hypo*, hypomethylated in activated preconditioned vs non-preconditioned; *Preconditioned-hyper*, hypermethylated in activated preconditioned vs non-preconditioned). The significance of enrichment is represented by the negative of the log of the p value, the enrichment fold change and the number of gene hits. **(G)** Transcription factor (TF) motif enrichment of a selected list of TFs (p value < 0.01) for each DMP cluster. The enrichment is represented by the negative of the log of the p value and the percentage of sequences matching the motif. **(H)** Boxplot representation of the beta values of four selected methylation sites across all studied groups. Each site is annotated according to its location in relation to the transcription start site (TSS) of the corresponding gene (+: downstream; -: upstream).

DNA methylation analysis of the four studied microglia conditions showed some differences (Figure 3E and Supplementary Table 3). When comparing non-activated microglia, both ATP-preconditioned and non-preconditioned, we identified hypomethylated DMPs (*ATP-hypo,* n=86) and hypermethylated DMPs (*ATP-hyper*, n=88) (p value < 0.05 and absolute mean Beta difference > 0.1). We also obtained both hypomethylated (*Preconditioned-hypo,* n=82) and hypermethylated (*Preconditioned-hyper,* n=111) DMPs when comparing activated ATP-preconditioned versus non-preconditioned microglia after LPS-mediated activation. GO analysis showed that both hypomethylation clusters are more enriched in inflammation-related terms; and the hypermethylated clusters (*ATP-Hyper* and *Preconditioned-Hyper*) are associated with neurotransmitter and purine-related terms. However, the level of enrichment significance was low, as demonstrated by the gene hits in each term (Figure 3F). TF motif enrichment analysis indicates high enrichment of Smad2 and Otx2 in *Preconditioned-Hypo* and *Preconditioned-Hyper*, respectively (Figure 3G). Looking into specific methylation sites showing differences in the preconditioning model, we identified, for instance, hypomethylation of *HLA-F* and *CD300LF* in both conditions exposed to ATP (activated and non-activated) and specific hypermethylation of *HPCA* in activated preconditioned microglia-like (Figure 3H).

ATP-preconditioning resulted in significant changes in gene expression. We detected 378 differentially expressed genes (DEGs) (205 upregulated and 173 downregulated) in ATP-preconditioned vs non-preconditioned non-activated microglia-like cells; and 1223 DEGs (667 upregulated and 556 downregulated) in ATP-preconditioned vs non-preconditioned microglia-like cells following LPS activation (Figure 4A). Of note, a significant proportion of the upregulated and downregulated DEGs were coincident in both comparisons (Figure 4B), suggesting that exposure to ATP by itself promotes gene expression changes that are maintained after activation. Unsupervised clustering of all DEGs from both comparisons separated them into five modules (M1-M5) (Figure 4C). From the GO analysis of all five modules, we focused on M1 and M3, both upregulated in activated preconditioned vs nonpreconditioned microglia-like cells. M1 is enriched in terms associated with phagocytosis, regulation of viral replication, IL-6, IL-10, TLR, and NF-kB signalling and antigen presentation (Figure 4D). M3 is enriched in cellular migration, and cytokine and chemokine activity (Figure 4D). Analysis of regulon enrichment based on gene expression of TF targets demonstrated a higher involvement of inflammation-related TFs, such as a member of the interferon-regulatory factor (IRF1), NF-kB subunits (RELA and NFKB1) and STAT proteins (STAT1 and STAT5B) (Figure 4E). To integrate the transcriptomics and DNA methylation data, we calculated the enrichment of the overlap between the genes from each gene expression module and the genes associated with the four DMP groups (*ATP-hypo, ATP-hyper, Preconditioned-hyper, Preconditioned-hypo*) (Figure 4F). Genes from M1 showed a higher overlap enrichment with the genes associated with *ATP-hypo* DMPs (demethylated CpG sites by ATP in non-activated cells). The overlap between M1 and *ATP-hypo* consisted in four CpG-gene pairs, that are demethylated after ATP stimulation, and upregulated in preconditioned microglia-like after the secondary LPS activation (Figure 4C and 4G). To investigate the potential acquisition of a trained or tolerized expression signature under our conditions, we used the lists of genes reported to be transcriptionally associated with trained immunity and immune tolerance in macrophages (31). In such study, six gene modules (Saeed-M1 - Saeed-M6) were defined, in which Saeed-M2 and Saeed M4 showed increased expression in trained macrophages, and Saeed-M3 and Saeed M5 were upregulated in tolerized macrophages. Here, we overlapped Saeed’s modules with the modules from the expression analysis of our model. The “Trained Immunity Signature” consisted of the genes associated with trained immunity in macrophages and upregulated in preconditioned microglia-like cells (M1-M3). The “Immune Tolerance Signature” consisted of the genes associated with tolerized macrophages and downregulated in preconditioned microglia-like cells (M4 and M5) (Figure 4H). The inspection of the expression of these signatures in activated preconditioned and non-preconditioned cells showed that ATP-treatment leads to the respective enhancement of trained immunity profiles and the hampering of tolerized profiles in microglia-like cells (Figure 4I).

**Figure 4.**
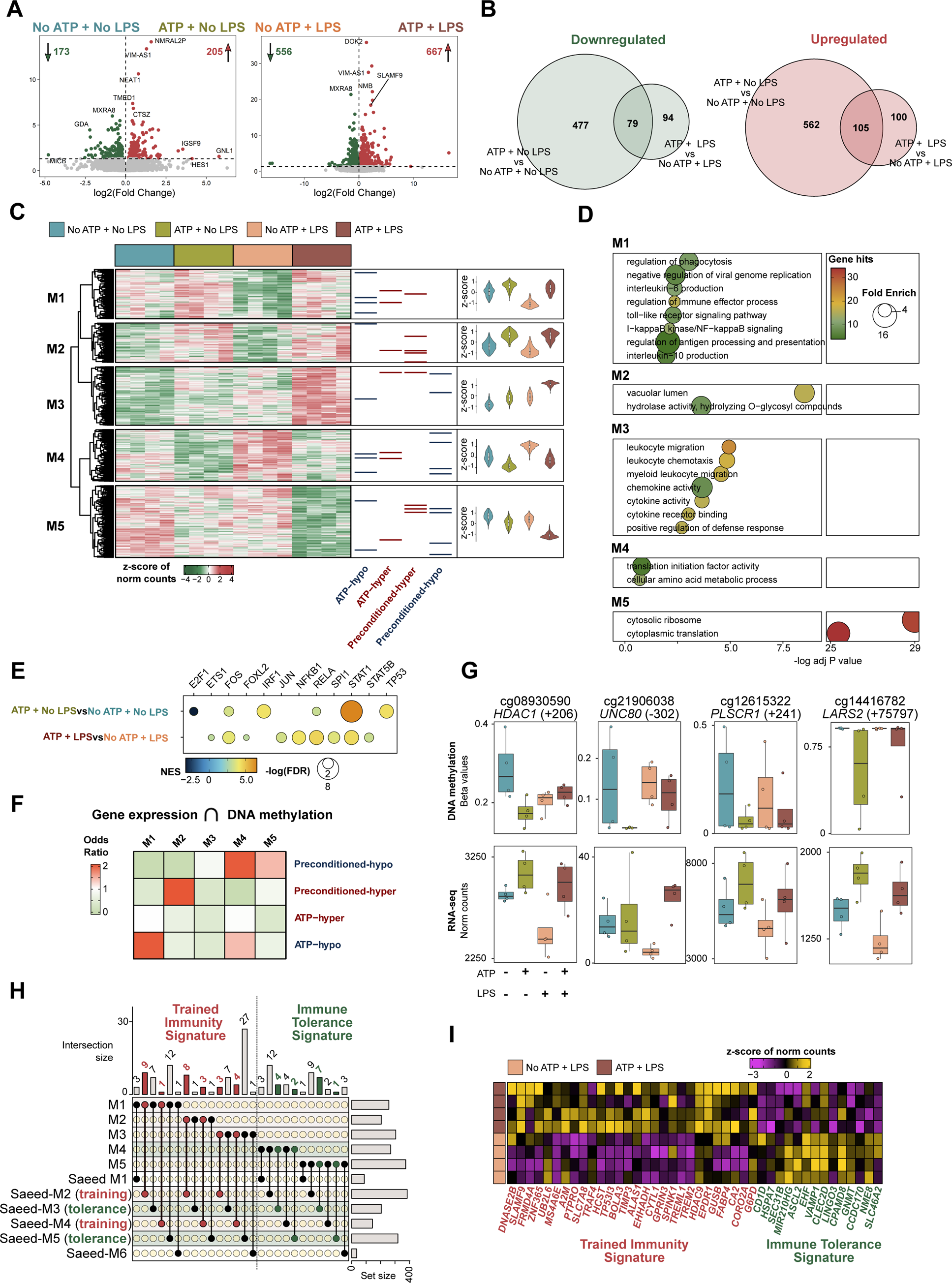
**(A)** Volcano plots depicting the differential expression between non-activated preconditioned and non-preconditioned microglia-like cells, and between activated preconditioned and non-preconditioned microglia-like cell. Differentially expressed genes (DEGs) are considered for FDR < 0.05. Upregulated DEGs (log2(Fold Change) > 0) are highlighted in red, and downregulated DEGs (log2(Fold Change) < 0) in green. **(B)** Venn diagram of the overlap between the list of genes up and downregulated in the two comparisons. **(C)** Heatmap representation of all the DEGs from both comparisons. Gene expression is represented as the z-score of the normalized counts. The significance cutoff was of FDR < 0.05. Unsupervised clustering divided the DEGs in five modules. DEGs associated with differentially methylated positions (DMPs) are highlighted (red: hypermethylated; blue: hypomethylated). The violin plots represent the mean z-score of each gene in each group. **(D)** Significantly enriched gene ontology (GO) terms for each of the lists of DEGs within each module. The significance of enrichment is represented by the negative of the log of the p value, the enrichment fold change and the number of gene hits. **(E)** Transcription factor (TFs) enrichment of all significant TFs (FDR < 0.05) in both differential expression comparisons (preconditioned vs non-preconditioned microglia-like cells before LPS; preconditioned vs non-preconditioned microglia-like cells after LPS). Significance is represented by the NES (Normalized Enrichment Score) and the negative of log of the adjusted p value (FDR). **(F)** Heatmap representation of the odds ratio of the overlap between the lists of the five DEG modules and the lists of genes associated with DMPs, obtained using *GeneOverlap*. Higher enrichment is represented by a higher odds ratio value (red). **(G)** Boxplot representation of the beta values (DNA methylation) and normalized counts (gene expression) of the four DMP-DEG pairs obtained from the overlap of M1 geness and hypomethylated DMPs in non-activated preconditioned vs non-preconditioned microglia-like cells. Each site is annotated according to its location in relation to the transcription start site (TSS) of the corresponding gene (+: downstream; -: upstream). **(H)** Overlap plot of the lists of genes from our gene expression modules (M1-M5) and the modules from Saeed *et al.* (31) (Saeed-M1 – Saeed-M6). Highlighted in red are the overlaps between the genes associated with ATP-preconditioning in our study (M1, M2, M3) and Saeed modules associated with immune training (Saeed-M2, Saeed-4). Highlighted in green are the overlaps between the genes associated with ATP-preconditioning in our study (M4, M5) and Saeed modules associated with immune tolerance (Saeed-M3, Saeed-5). We highlighted 26 genes deemed as the “Immune Training Signature”, and 14 genes deemed as the “Immune Tolerance Signature”. **(I)** Heatmap representation of the genes from the “Immune Training Signature” and the “Immune Tolerant Signature” in microglia-like cells activated non-preconditioned and preconditioned. Gene expression is represented as the z-score of the normalized counts.

### ATP-mediated preconditioning induces a molecular signature in microglia-like cells that correspond to disease inflammatory macrophage subsets expanded in epilepsy

Finally, we examined the transcriptomic changes observed in ATP preconditioned microglia-like cells in the context of human microglia subpopulations in patients with epilepsy and healthy individuals. To this end, we used a snRNA-seq dataset consisting of 24,037 microglia nuclei from both patients with epilepsy and healthy controls without any diagnosed neuropathology. The integrated object consisted of nine clusters of microglial nuclei, expressing high levels of canonical markers, *P2RY12* and *CX3CR1,* and obtained after exclusion of nuclei expressing markers for macrophages, T cells and other CNS cell types (Figure 5A). Each subpopulation showed variable expression of known homeostatic genes (*P2RY12* and *CX3CR1*) and pathology-related microglia markers (*SPP1* and *APOE*). In addition, we investigated the average expression of gene signatures characterizing the phenotypes of “disease-associate microglia” (DAM) and “disease-inflammatory macrophages” (DIMs), described elsewhere (32) (Figure 4I). We observed that the genes from M1 (upregulated in activated preconditioned vs nonpreconditioned microglia-like cells described in our microglia-like RNA-seq analysis, presented high expression in cluster 3, whose transcriptomic profile corresponded to the one from DIMs (Figure 5B). We then subsetted the snRNA-seq object, considering only for cluster 3, which consisted of 532 and 3155 nuclei from epilepsy patients and controls, respectively. Reclustering of these nuclei resulted in five subclusters, two of which (subclusters 0 and 4) are expanded in epilepsy in comparison to controls (Figure 5C and 5D). Furthermore, the expression of M1 genes and the “Trained Immunity Training Signature” were respectively increased in subclusters 0 and 4 (Figure 5E), both expanded in samples from epilepsy patients, supporting that ATP preconditioning induces an upregulation signature associated with epilepsy.

**Figure 5.**
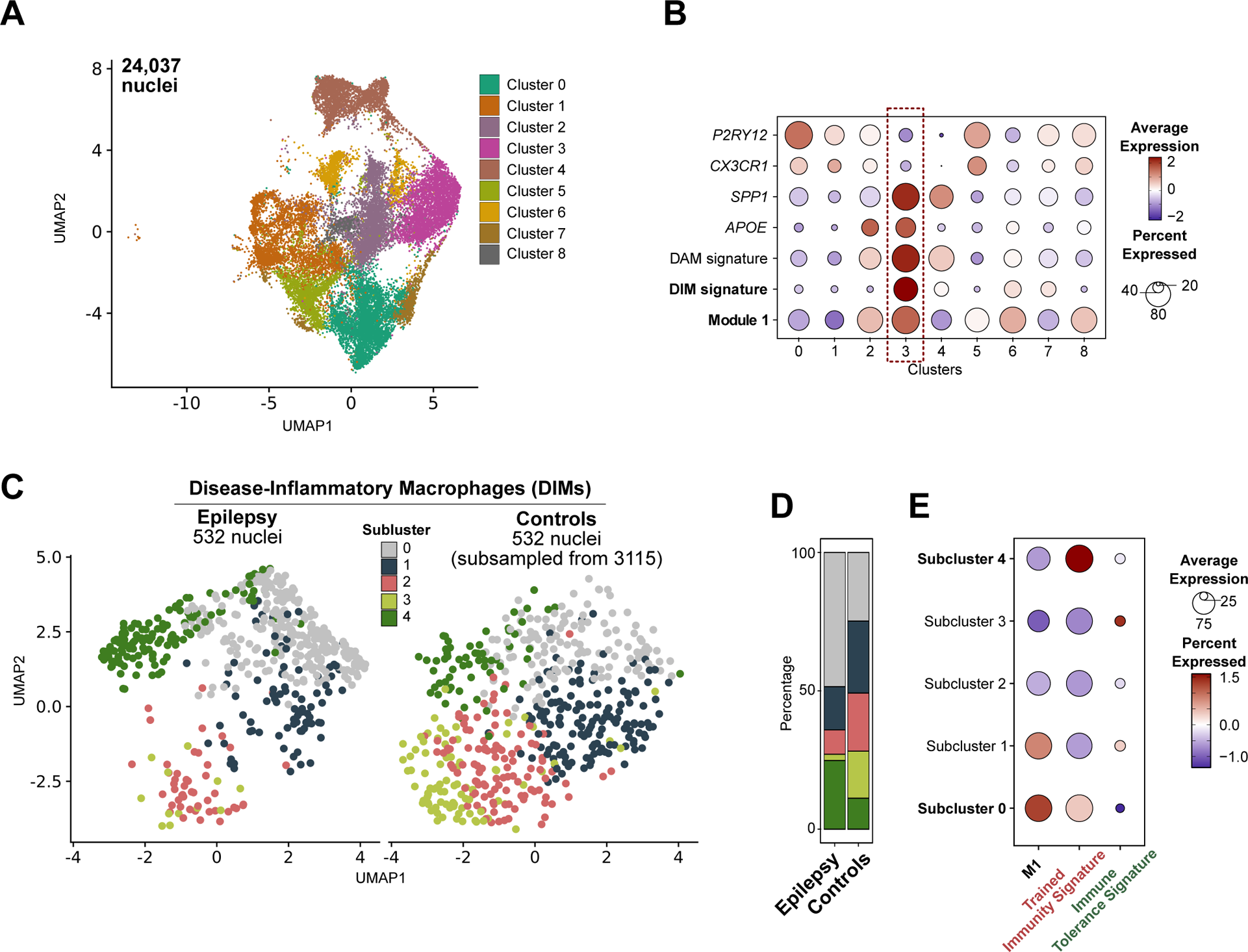
**(A)** UMAP representation of the integrated single-nucleus (sn)RNA-seq object composed of 24,037 microglia nuclei from epilepsy patients and healthy individuals, distributed across nine clusters **(B)** Dot plot representation of the expression of genes associated with homeostatic (*P2RY12* and *CX3CR1*) and pathological (*SPP1* and *APOE*) microglial phenotypes, and the Module Score expression of gene signatures characteristic of “disease-associated microglia” (DAM) and “disease inflammatory-macrophages” (DIMs), and the M1 genes from our preconditioning study. **(C)** UMAP representation, split by pathological conditions (*Epilepsy* and *Controls*), of only the nuclei from cluster 3. Reclustering of the DIM population resulted in five subclusters (cluster 0-4). The object accounts for 532 nuclei from epilepsy patients and 3155 nuclei from controls. For representation purpose, only 532 randomly selected nuclei from controls (representative of the whole population) were presented. **(D)** Proportion distribution of the five DIM subpopulations in epilepsy patients and controls. Cluster 0 and 4 are expanded in epilepsy in comparison to controls. **(E)** Dot plot representation of the expression of the Module Score of the genes from M1, the “Immune Training Signature” and the “Immune Tolerant Signature”.

## DISCUSSION

In this study, we have shown that ATP-mediated preconditioning of microglia-like cells is accompanied by epigenetic and transcriptional reprogramming that is associated with their subsequent inflammatory responses. The occurrence of innate immune memory and the participation of epigenetic changes in this context provides new potential mechanisms to explain some of the roles of microglia in neuropathology. Secondary immune responses following a first insult can exhibit different outcomes depending on the nature of the stimuli. In the case of trained immunity, the response is more pronounced or exacerbated, whereas in the case of tolerance, the response is diminished (8). Evidence supporting the existence of memory in microglia remains limited. Nevertheless, studies have shown that sequential treatment of microglia with LPS can influence the production of cytokines and reactive oxygen species, phagocytic activity and persistent alterations in gene expression profiles (9,10). Processes leading to the acquisition of innate immune memory are thought to be tightly regulated by epigenetic modifications. Two studies have conducted RNA-seq, ATAC-seq and ChIP-seq of multiple histone marks on microglia, revealing epigenetically-dependent transcriptomic changes associated with immune preconditioning (11,12). The participation of DNA methylation was not explored in these studies, despite its relevance for myeloid differentiation. DNA methylation is the most stable and one of the best studied epigenetic modifications. It consists of the addition of a methyl group to cytosine nucleotides in the DNA and contributes and/or reflects the transcriptional competence of chromosomal regions (33,34)

As mentioned above, the use of human microglia models is necessary to study the molecular mechanisms involved in pathogenesis. A common concern regarding the use of microglia-like models is their distinct ontogeny compared to *in vivo* microglia, which differentiate from precursor cells that migrate into the brain in early embryogenesis and are independent from peripheral monocytes (35,36). A common *in vitro* approach to study human microglia is iPSC-derived microglia, which aims at emulating the microglial ontogeny (37). Using microglia-like cells derived from monocytes offers the advantage of a simpler and quicker experimental protocol in comparison to iPSC-derived microglia, which consists of longer multi-step procedure (38). Furthermore, the use of the differentiation of microglia-like cells from monocytes has gained momentum due to a recent discovery of a novel myeloid population within the cerebral microglial population. Using single-cell transcriptomics, Silvin and colleagues (2022), described a subset of cells known as “disease-inflammatory macrophages” (DIMs). Although DIMs share features with microglia, what made being annotated under the microglia spectrum, they are actually derived from infiltrating monocytes. These DIMs present a general pro-inflammatory profile with neurotoxic implications in neurodegeneration and aging (32).

*In vitro* models of microglia-like cells have been shown to mimic *in vivo* microglia features, including morphology, specific surface protein expression, expression of specific canonical homeostatic markers and overall transcriptomic profile (25,27,28,38–40). In this study, we have performed a multi-level characterization of microglia-like cells differentiated from monocytes demonstrating the acquisition of the elongated and ramified structure, and the acquisition of standard microglia surface markers. At the transcriptomic level, our microglia-like cells showed high resemblance with primary human microglia, outperforming those generated by previous protocols. Furthermore, we obtained the DNA methylation profiles of microglia-like cells, which also support its cellular identity, given their cell type specificity (41). In this analysis, we have demonstrated a successful replication of microglial DNA methylation pattern. The DNA methylation profiles of microglia-like cells cluster closer to *in vivo* primary microglia than to primary monocytes and are enriched for TF binding motifs associated not only with myeloid cells but also with microglia. In an encompassing study, Gosselin *et al.* performed transcriptomic and epigenomic characterization of mice and human microglia, including ATAC-seq (defines open chromatin), and ChIP-seq for H3K4me2 (demethylation of histone H3 lysine 4, annotates regulatory regions like enhancers) and H3K27ac (acetylation of histone H3 lysine 27, annotates regions with high transcriptional activity). ATAC-seq peaks associated with H3K4me2 and H3K27ac in microglia were predominantly enriched for PU.1, denoted as the main driver of microglial identity, but also for IRF, MEF2, SMAD and MAF factors. Our results indicate that these same factors, with prominence for PU.1, are associated with the DNA methylation changes in microglia-like differentiation.

Our model of microglia preconditioning aligns with the pathogenesis landscape observed in various neurodegenerative diseases, characterized by the presence of widespread primed/activated microglia states reported in these conditions (8). In addition, we recognize the specific relevance of integrating purinergic signalling within this framework, particularly when considering its implications in the context of epileptogenesis. Mesial temporal lobe epilepsy with hippocampal sclerosis (MTLE-HS) is particularly relevant considering the high incidence of reported initial precipitating injuries in early childhood, namely trauma, hypoxia, intracranial infection and febrile seizures (42–45). It could be postulated that purine-driven microglia priming caused by the initial insult may influence posterior microglia responses in ways that could influence the structural and molecular changes associated with the latent phase of epilepsy development. Furthermore, considering the progressive nature of MTLE, seizure activity may precondition microglia activation through the release of extracellular ATP and adenosine, with impact on the continuous aggravation of neuroinflammation and prognosis worsening. An important aspect that deserves attention in this discussion is the previously described adenosine kinase (ADK) hypothesis of epileptogenesis, which proposes that, in epileptogenic conditions, the levels of ADK, an enzyme responsible for converting endogenous adenosine into AMP, increase. This elevation of ADK results in the extracellular depletion of adenosine (46). This mechanism is directly associated with DNA methylation. The conversion of S-adenosylmethionine (SAM) to S-adenosylhomocysteine (SAH) serves as a major source of methyl groups. Subsequently, SAH is hydrolysed to produce homocysteine and adenosine. Hence, any disturbance in the intracellular and extracellular balance of adenosine levels can directly impact DNA methylation by altering the thermodynamics of the SAM metabolism. In the early stages, this process was proposed to be unidirectional leading to overall DNA hypermethylation due to increased adenosine consumption by ADK in the context of epileptogenesis (47,48). However, studies have demonstrated that in complex and heterogeneous pathologies such as epilepsy, DNA methylation alterations occur in a region-specific manner, with some methylation sites showing hypermethylation and others hypomethylation. In a recent study conducted by our group, we observed that DNA methylation of multiple sites associated with neuroinflammation correlate with disease duration, suggesting the implication of microglia in epilepsy progression (49).

In the present study, we have observed that ATP-driven preconditioning is associated with the acquisition of a more pro-inflammatory phenotype. This shift in the phenotype is accompanied by bidirectional changes in both transcriptomic and DNA methylation profiles, with a tendency towards DNA hypomethylation in pathways associated with inflammation. Furthermore, activated preconditioned microglia-like cells acquired an immune training gene expression profile similar to the previously described in macrophages treated with β-glucan (31). Furthermore, we were able to replicate the changes caused by ATP preconditioning *in vitro* in snRNA-seq data from epilepsy patients and controls. These results provide support for the replicability of these mechanisms described in the context of epileptogenesis. Additionally, we have determined that genes upregulated in activated and preconditioned microglia-like cells, as well as the genes associated with immune training, are elevated in subpopulations of DIMs that are expanded in epilepsy (Figure 6).

**Figure 6.**
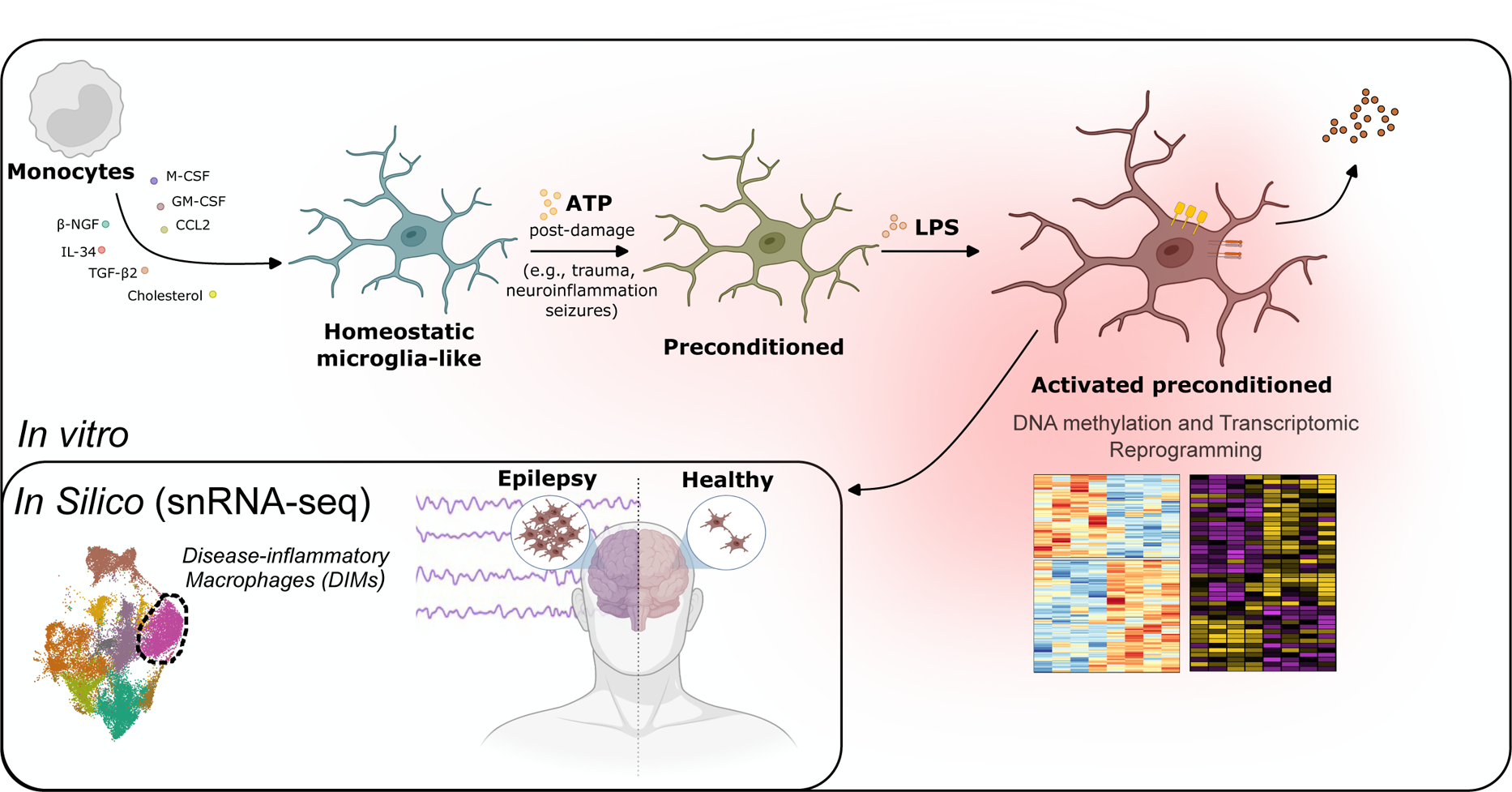
Schematic illustration of the summarized results from this study. ATP-preconditioning of microglia-like cells, mimicking post-damage the increase in purine extracellular levels after damaging events (including trauma, neuroinflammation and epileptic seizures), promotes an exacerbated pro-inflammatory response upon activation, which is coupled with DNA methylation modifications and a transcriptomic shift towards an immune training phenotype. This pro-inflammatory trained profile is enriched in subpopulations of “Disease-inflammatory Macrophages” (DIMs) that are expanded in snRNA-seq data from epilepsy patients in comparison to healthy individuals.

Our *in vitro* model utilizing microglia-like cells has demonstrated that exposure to ATP, which mimics the conditions observed during brain insults such as epileptic seizures, leads to a concurrent increase in inflammatory activation and alterations in DNA methylation and transcriptional patterns. These changes encompass functional categories and genes that support the occurrence of epigenetic priming. Various studies have used similar models in patients to study pathological mechanisms related to microglia (25,50–54). By comparing our ATP-mediated preconditioning model with microglia subsets from snRNA-seq data derived from epilepsy patients and healthy donors, we provide evidence for the involvement of purinergic conditioning in the acquisition of disease associated microglial states. Activated preconditioned microglia-like cells resemble subpopulations of DIMs that are expanded in the brains of epilepsy patients, potentially contributing to the heightened neuroinflammatory profile associated with epileptogenesis. Further epigenetic and functional analysis of microglia cells from patients with MTLE-HS compared to controls will shed light on the distinct responsiveness of microglia in epilepsy.

## MATERIALS AND METHODS

### Monocyte purification and in vitro differentiation to microglia-like cells

Peripheral blood was collected under the form of buffy coats obtained from anonymous donors through the Catalan Blood and Tissue Bank. All donors signed an informed consent and sample collections were performed as per the guidelines of the World Medical Association (WMA) Declaration of Helsinki.

Peripheral blood mononuclear cells (PBMCs) were isolated using Ficoll-Paque gradient centrifugation. Monocytes were then purified using positive selection with CD14+ MicroBeads (Miltenyi Biotec). For microglia-like differentiation, monocytes were cultured in RPMI Medium 1640 + GlutaMAX^TM^ (Gibco, Thermo Fisher) containing 100 units/mL penicillin, 100 µg/mL streptomycin, and supplemented with 10 ng/mL human M-CSF, 10 ng/mL human GM-CSF, 10 ng/mL human β-NGF, 100 ng/mL human IL34, 100 ng/mL human CCL2, 2 ng/mL human TGF-β2 (PeproTech) and 1.5 µg/mL cholesterol solution dissolved in ethanol. Microglia-like plating density was of 1.5 M cells per well (6-well plates). Homeostatic microglia-like cells were obtained after seven days of culture without media changes. For monocyte-derived macrophage differentiation, monocytes were attached to plates by incubation with serum-free medium and posteriorly cultured in α-minimal essential medium (α-MEM; Invitrogen, Carlsbad, CA, USA) containing 10% foetal bovine serum, 100 units/mL penicillin, 100 µg/mL streptomycin and supplemented with 25 ng/mL human M-CSF. Plating density for monocyte-derived macrophages was 3 M cells per well (6-well plates). No media changes were performed for the macrophage culture, and cells were activated with 10 ng/mL of LPS at day six of culture and collected 24h after.

### Activation and preconditioning of microglia-like cells

The design of the preconditioning analysis consisted in the development of the following four microglia-like activation states: microglia-like without any stimuli and collected at day seven (non-activated non-preconditioned); microglia-like stimulated with 100 µM ATP at day six and collected 24h after (non-activated preconditioned); microglia-like stimulated with 100 ng/mL LPS at day eight and collected 48h after (activated non-preconditioned); Microglia-like stimulated with 100 µM ATP at day six and with 100 ng/mL LPS at day eight, and collected at day ten of culture (activated preconditioned).

### Real-time quantitative reverse-transcribed polymerase chain reaction (qRT-PCR)

Total RNA was isolated using the Maxwell RSC simplyRNA Cells Kit (Promega) and reverse-transcribed to cDNA with Transcriptor First Strand cDNA Synthesis Kit (Roche) per the manufacturer’s instructions. qRT-PCR primers were designed with Primer3 software (55) (*P2RY12*: forward – CCACTCTGCAGGTTGCAATA; reverse – GGCTTGCATTTCTTGTTGGT; *CX3CR1*: forward – CACAAAGGAGCAGGCATGGAAG; reverse – CAGGTTCTCTGTAGACACAAGGC; *RPL38* (housekeeping): forward – TGGGTGAGAAAGGTCCTGGTC; reverse - CGTCGGGCTGTGAGCAGGAA. Technical triplicates were run for each sample using LightCycler® 480 SYBR Green Mix (Roche) and analysed with a LightCycler instrument (Roche).

### RNA-seq

RNA-seq libraries of monocytes and microglia-like cells were generated and sequenced by Novogene (Cambridge), in 150-bp paired-end, with the Illumina NovaSeq 6000 platform, using four biological replicates for each group. An average of more than 65 million reads were obtained for all samples. Fastq files were align to the hg19 transcriptome using *HISAT2* (56) with default settings, and read counts by gene were assigned with *featureCounts* (57). All the posterior analysis was performed using R language. Differentially expressed genes (DEGs) were calculated using *DESeq2* (58). Donor was used as a covariate in the model. Significant DEGs were considered for FDR < 0.05. Variance Stabilizing Transformation (VST) and normalized count values were used for representation.

For the integration with public RNA-seq, raw count values were merged and adjusted for dataset using the *ComBat* function of the *sva* package (59). Raw count data for primary adult and foetal human microglia, C20 and HMC3 cell lines, CD14 and CD16 monocytes, dendritic cells, monocyte-derived macrophages, monocyte-derived microglia-like cells, and iPSC-derived microglia was collected from Rai *et al* (2020) (28). This study included data from other public datasets, namely GSE89189 and GSE117829.

### DNA methylation profiling, data processing and analysis

Genomic DNA was isolated from cell lysates in Proteinase K using an in-house salting out protocol. We then performed bisulfite conversion with the EZ DNA Methylation-Gold^TM^ Kit (Zymo Research, Irvine, CA, USA) and DNA methylation profiling using Infinium MethylationEPIC BeadChips. These arrays cover 850,000 single-nucleotide positions, accounting for 99% of the annotated RefSeq genes. Methylation data was pre-processed and analysed using R language with the *shinyÉPICo* web interface (60) base on the *minfi* (61) and *limma* (62) packages. Beta values, comprised between 0 and 1 (0 and 100 % ratio of methylated probe intensity/sum of methylated and unmethylated probe intensities), were used for visualization purposes. M values consist of the log2 ratio of the intensities of the methylated and unmethylated probes and were used for statistical purposes, since beta values are heteroscedastic for highly methylated and unmethylated sites (63). Normalization was performed by the Noob and Quantile methods from *minfi*. Probes with a detection significance of p < 0.01 were removed. CpHs positions were kept in the dataset. Since all subjects were male, methylation sites from X and Y chromosomes were kept, whereas all positions located at single nucleotide polymorphisms (SNP) loci were removed (minimum allele frequency (MAFs) = 0). Differentially methylated positions (DMPs) were calculated using an eBayes-moderated t test from *limma*. Public DNA methylation data of primary microglia (n=56) isolated from postmortem brain samples (GSE191200) was integrated in the in-house generated data, pre-processed, and normalized all together and batch corrected with the *ComBat* function from the *sva* package (59).

### Quantification of supernatant cytokines

Supernatants from cell cultures were collected and stored at −80 °C. Before use, frozen samples were thawed at room temperature. Enzyme-linked immunosorbent assays (ELISA) were performed to detect IL-1β (Invitrogen, Thermo Fisher Scientific).

### Flow cytometry

Levels of cell surface protein markers were evaluated by flow cytometry using a BD LSR Fortessa™ cytometer. Cells were washed once with PBS, after which Versene, a non-enzymatic dissociation buffer (ThermoFisher), was added for detachment. After the addition of RPMI Medium 1640 + GlutaMAXTM (Gibco, Thermo Fisher) containing 100 units/mL penicillin, 100 µg/mL streptomycin and 10 % FBS to the wells, cells were gently scrapped and collected. After collection, cells were resuspended in the staining buffer (PBS with 4% FBS and 2 mM EDTA). Cells were incubated at 4°C with the viability dye LIVE/DEAD^TM^ Fixable Violet (ThermoFisher), following the manufacturer’s protocol. Cell staining followed a protocol developed in-house consisting of two separate panels of antibodies [Panel 1: CD14 (FITC) (#367116 BioLegend), CX3CR1 (PE) (#341604 BioLegend), CD11b (PE-Cy5) (#301308 BioLegend), HLA-DR (PE-Cy7) (#307616 BioLegend), CD192 (APC) (#357208 BioLegend), CD68 (APC-Cy7) (#333822 BioLegend), CD45 (KO) (#B36294 BeckmanCoulter Life Sciences); Panel 2: CD14 (FITC), P2RY12 (PE) (#392104 BioLegend), CD11b (PE-Cy5), HLA-DR (PE-Cy7), MERTK (APC) (#367612 BioLegend), CD64 (APC-Cy7) (#305026 BioLegend), CD45 (KO)]. After staining, cells were fixed in PBS + 4% paraformaldehyde and analysed within the following 24 hours. A compensation matrix was developed from single-positive experiments with beads. Median fluorescence intensity (MFI) values were calculated by subtracting the value from the unstained control to the stained sample.

### Pre-processing, integration, and analysis of single-nucleus (sn)RNA-seq data

We used public snRNA-seq data from brain tissue samples of surgically treated epilepsy patients and autopsied individuals (64–67). The data was obtained in the count matrix format, at the raw or pre-processed stages. Demultiplexing had already been done and the sequences had been aligned to the human reference genome (GRCh38), accounting for intronic and exonic regions. The individual samples in each dataset were concatenated into a single count matrix, and the subsequent processing was performed using the *Seurat* (*v4.0.2*) R package (68). Genes were only considered if detected in at least three nuclei, and nuclei were excluded if presented unique genes inferior to 200 or superior to 5,000, total UMI counts less than 500 and over 20,000, mitochondrial RNA content superior to 20% and ribosomal RNA content superior to 5%. In addition, a list of 105 genes shown to be influenced by postmortem interval in cerebral cortex were filtered out (69). Potential doublets were estimated using the *doubletFinder_v3* function (70) for each individual subject and removed. The individual Seurat objects for each dataset were normalized using *SCTransform* normalization (71) with default 3000 variable genes. Dimensionality reduction was performed with Principal Component Analysis (PCA) and Uniform Manifold Approximation and Projection (UMAP) accounting for the 30 main principal components (PCs). The annotation for the main cell types (neurons, oligodendrocytes, OPCs, astrocytes, endothelial cells, and immune cells) in the CNS was performed for each dataset using the *FindNeighbors* (30 PCs) and *FindClusters* (res = 0.05) functions, after which known canonical gene markers were identified within the lists of cluster DEGs obtained using FindAllMarkers (min.pct = 0.1 and logfc.threshold = 0.25, test.use=Wilcox). The immune cell clusters were annotated based on high expression of *CD74, DOCK8, APBB1IP, HLA-DRA, PTPRC, P2RY12, C1QB, CX3CR1, C3, CSF1R* and *AIF1*. Individual subjects with unproportionally low nuclei number in relation to the other subjects in each dataset were removed before integration. The immune cell clusters from all datasets were integrated using the Seurat pipeline (72) adapted for *SCTransform* normalization. The main 20 PCs and the 3000 most variable features across all datasets were considered for the *FindIntegrationAnchors* and *IntegrateData* functions. Dimensionality reduction was performed using PCA and UMAP (50 PCs). Clustering was performed using *FindNeighbors* (50 PCs) and *FindClusters* (res = 0.25). Clusters corresponding to macrophages (based on *MRC1* expression), T cells (based on *CD247* expression) and potential doublets (based on *SLC1A2, PLP1* and *VCAN* expression) were removed, resulting in a final integrated object with 24,037 microglia nuclei grouped into nine populations. For the visualization of gene expression, we used the normalized and scaled counts of the “RNA” assay. The average expression of the lists of geneswas obtained with *AddModuleScore*.

### Statistics, data analysis and representation

The R 4.2.0. software was used for statistical analyses. Group medians were compared using the Mann-Whitney test for numeric variables. Fisher’s exact test was used to calculate the significance of non-random association between two categorical variables. Heatmaps were developed with *heatmap.2* function of the *gplots* package (73), for z-score estimation, followed by the *Heatmap* function of the *ComplexHeatmap* package (74). The representation of DNA methylation in relation to genomic coordinates was obtained using the *Gviz package* (75). Overlaps were obtained using either the *VennDiagram* package (76) or the *<u>UpSet</u>*function from *ComplexHeatmap*. Transcription factor (TF) motif enrichment for DMPs was obtained using the *findMotifsGenome.pl* function of HOMER (Hypergeometric Optimization of Motif EnRichment) (77), considering a window of ± 250 bp. Gene ontology (GO) enrichment for DMPs and determination of CpG-gene pairs were performed using the GREAT online tool (http://great.stanford.edu/public/html) (78), with default settings. All EPIC array coordinates were used as background for motif enrichment and GO analyses of methylation data. Functional enrichment of differential expression data from the RNA-seq was performed using the Discriminant Regulon Expression Analysis (DoRothEA) (79) for TF activity, and the *enrichGO* function from the *clusterProfiler* package (80), for GO. DNA methylation and RNA-seq results were integrated by evaluating the enrichment of the overlap between DEGs and DMP-related genes using a Fisher’s exact test within *GeneOverlap* (81) and Gene Set Enrichment Analysis (GSEA) using *fgsea* (82).

## ACKNOWLEDGEMENTS

We thank the CERCA Programme/Generalitat de Catalunya, the Josep Carreras Foundation and ICBAS-UP for institutional support. EB is funded by the Spanish Ministry of Science and Innovation (MICINN) [PID2020117212RB-I00; AEI/10.13039/501100011033]. RM-F is funded by an FCT (*Fundação para a Ciência e Tecnologia*) fellowship (SFRH/BD/137900/2018). UMIB is funded by FCT Portugal (UIDB/00215/2020 and UIDP/00215/2020), and ITR (LA/P/006/2020).

We thank all members of the Epigenetics and Immune Disease Group at the Josep Carreras Leukaemia Research institute, the Immunogenetics Laboratory of the Molecular Pathology and Immunology Department of the ICBAS-UP, and the Neurophysiology and Neurology Departments of *Centro Hospitalar Universitário de Santo António* (CHUdSA).

## AUTHORS’ CONTRIBUTIONS

R.M.-F., B.L. and E.B. conceived the study. R.M.-F., J.C.-S. and L.C. contributed to laboratory work. R.M.-F. and J.C.-S. performed bioinformatic analysis. R.M.-F., J.C.-S., B.L. and E.B. contributed to the verification and interpretation of the analysis, and to the writing of the article. All the remaining authors reviewed, edited and approved the final version of the manuscript.

