## Supplementary Materials for "Purinergic Preconditioning Induces Epigenomic and Transcriptomic Changes Resembling Epilepsy-associated Microglial States"

### Supplementary Figure 1

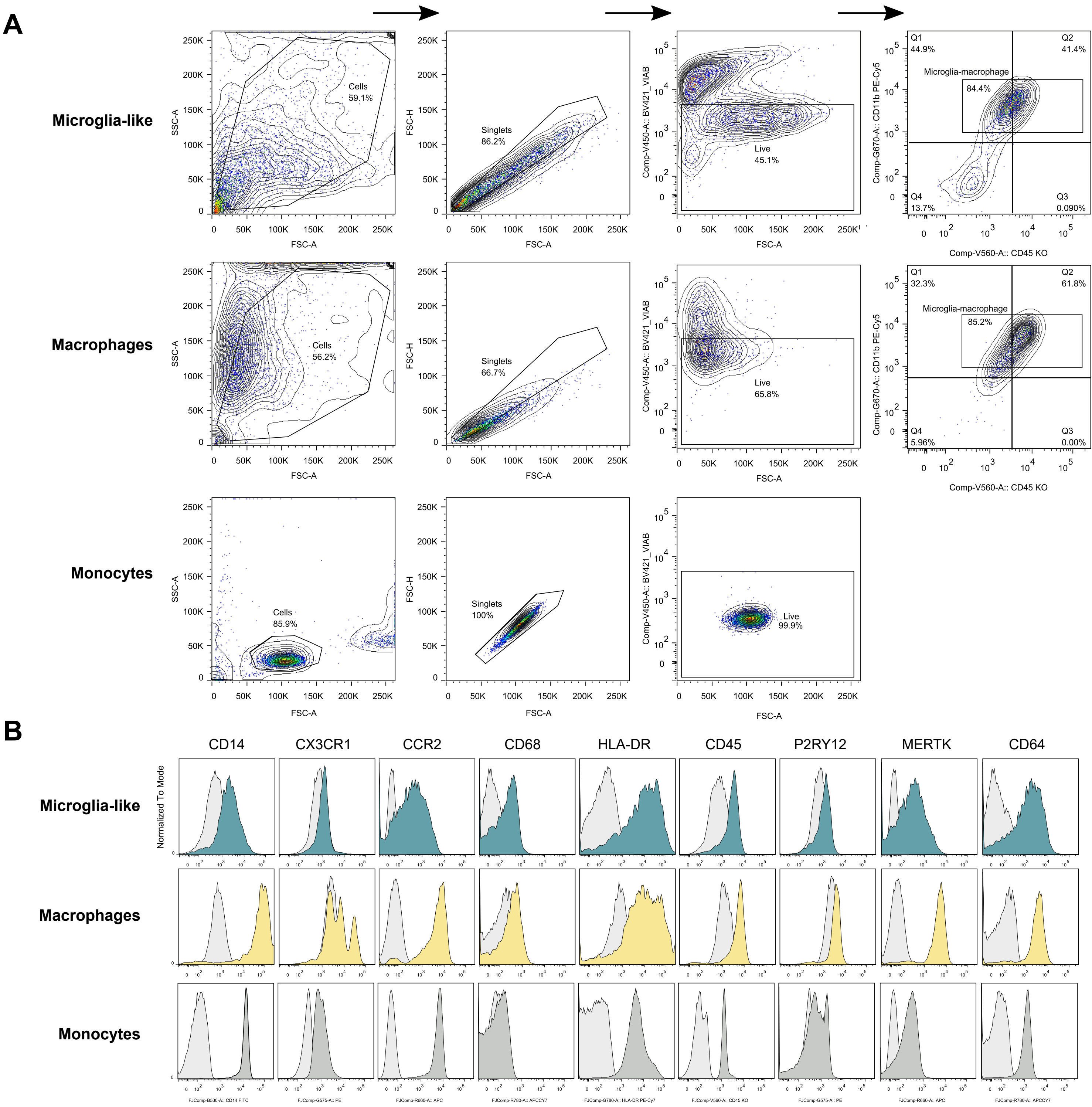

### Supplementary Figure 2

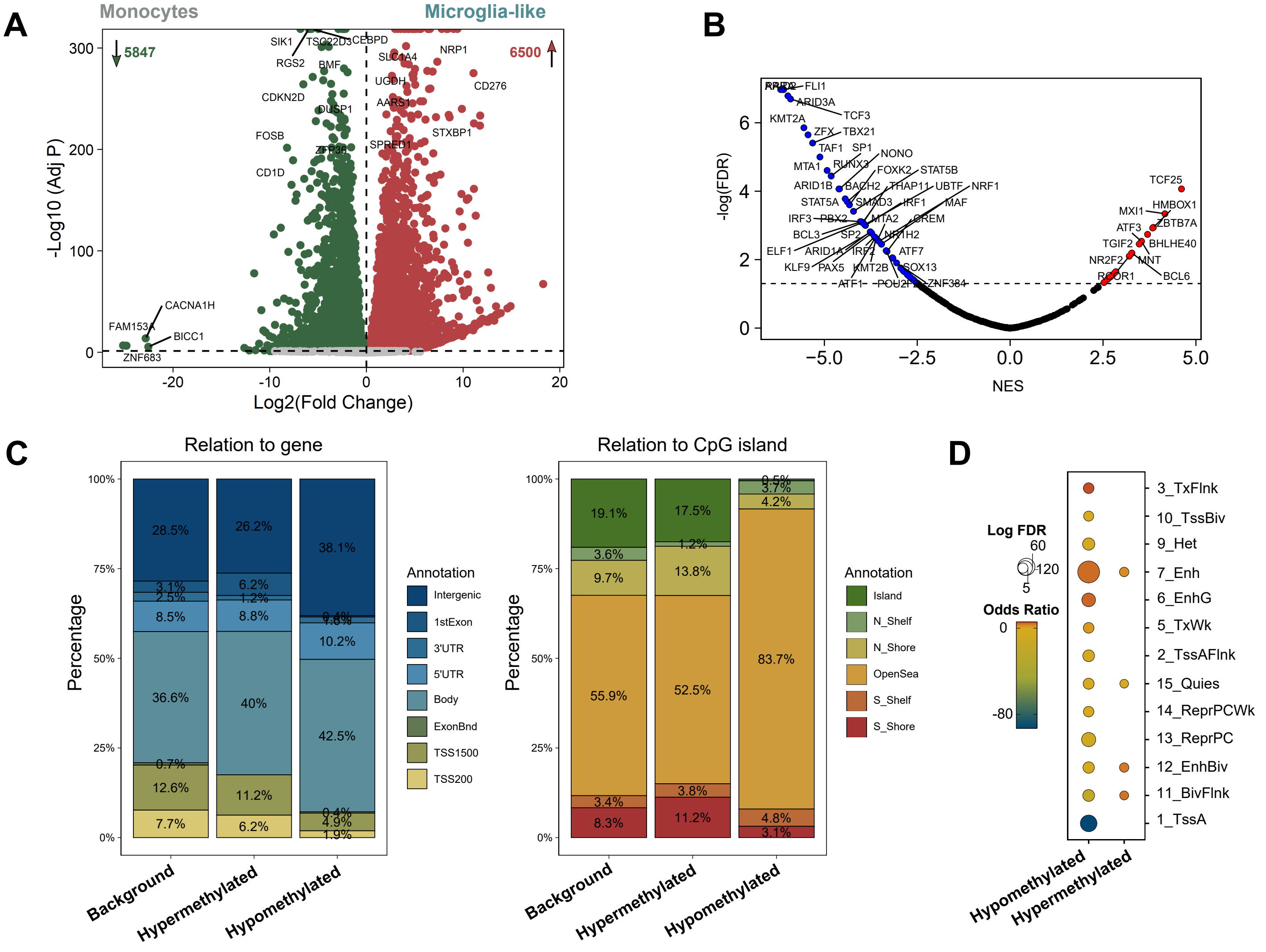

### Supplementary Figure 3

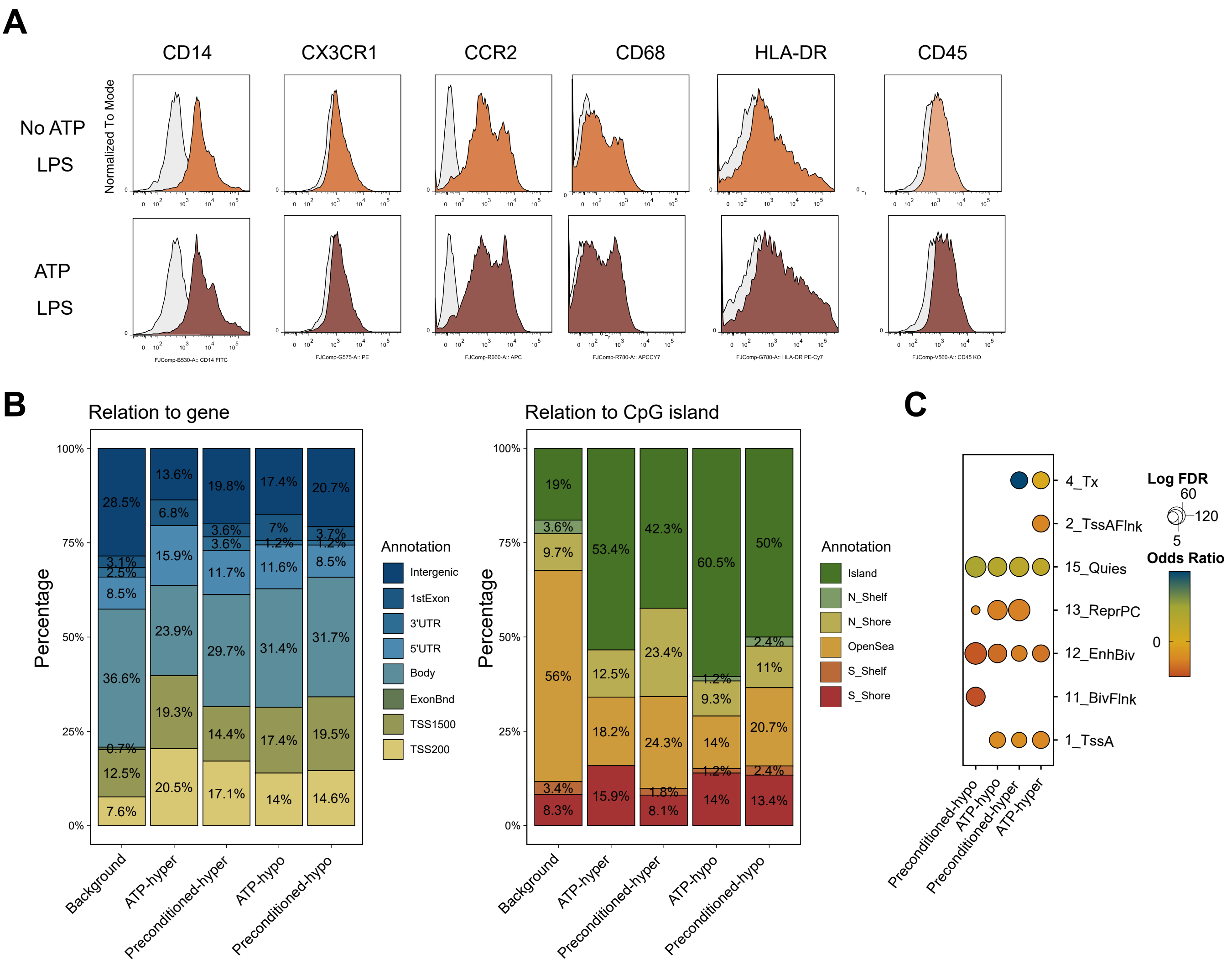

#### SUPPLEMENTARY FIGURE LEGENDS

**Supplementary Figure 1. (A)** Representation of the flow cytometry gating strategy used to single out the population of microglia/macrophage populations in the *in vitro* models of monocyte-derived microglia-like and monocyte-derived macrophages. Microglia and macrophages are CD11b+, and present low or high CD45 surface levels, respectively. For monocytes, we considered all live cells. **(B)** Histogram representation of the protein levels of CD14, CX3CR1, CCR2, CD68, HLA-DR, CD45, P2RY12, MERTK and CD64 obtained for freshly isolated monocytes, microglia-like and macrophages. **(C)** Proportion distribution of the lists of hypermethylated and hypomethylated differentially methylated positions (DMPs) in microglia-like vs monocytes in relation to gene (left panel) and in CpG islands (right panel), according to annotations from the Infinium MethylationEPIC array. The proportion of each category was calculated for background, hypermethylated and hypomethylated DMPs. N, north; S, south; 1stExon, first exon; UTR, untranslated region; ExonBnd, exon boundary; TSS, transcription start site. **(D)** Enrichment of hypermethylated and hypomethylated DMPs in microglia-like vs monocytes in ChromHMM categories of monocytes (Roadmap Epigenomics Project). Fisher's exact tests were calculated using all the positions annotated in the EPIC array as background.

**Supplementary Figure 2. (A)** Volcano plot depicting the differential expression between microglia-like cells and monocytes. Differentially expressed genes (DEGs) are considered for FDR < 0.05. Upregulated DEGs (log2(Fold Change) > 0) are highlighted in red, and downregulated DEGs (log2(Fold Change) < 0) in green. **(B)** Transcription factor (TFs) enrichment of all significant TFs (FDR < 0.05) in the differential expression comparison between microglia-like cells and monocytes. Significance is represented by the NES (Normalized Enrichment Score) and the negative of log of the adjusted p value (FDR). **(C)** Proportion distribution of the lists of hypermethylated and hypomethylated differentially methylated positions (DMPs) in microglia-like vs monocytes in relation to gene (left panel) and in CpG islands (right panel), according to annotations from the Infinium MethylationEPIC array. The proportion of each category was calculated for background, hypermethylated and hypomethylated DMPs. N, north; S, south; 1stExon, first exon; UTR, untranslated region; ExonBnd, exon boundary; TSS, transcription start site. **(D)** Enrichment of hypermethylated and hypomethylated DMPs in microglia-like vs monocytes in ChromHMM categories of monocytes (Roadmap Epigenomics Project). Fisher's exact tests were calculated using all the positions annotated in the EPIC array as background.

**Supplementary Figure 3. (A)** Histogram representation of the protein levels of CD14, CX3CR1, CCR2, CD68, HLA-DR and CD45 obtained for freshly isolated monocytes, and microglia-like preconditioned and non-preconditioned after LPS stimulation. **(C)** Proportion distribution of the lists of hypermethylated and hypomethylated differentially methylated positions (DMPs) in non-activated preconditioned vs non-preconditioned (*ATP-hyper* and *ATP-hypo*) and activated preconditioned vs non-preconditioned (*Preconditioned-hypo* and *Preconditioned-hyper*) in relation to gene (left panel) and in CpG islands (right panel), according to annotations from the Infinium MethylationEPIC array. The proportion of each category was calculated for background, hypermethylated and hypomethylated DMPs. N, north; S, south; 1stExon, first exon; UTR, untranslated region; ExonBnd, exon boundary; TSS, transcription start site. **(D)** Enrichment of hypermethylated and hypomethylated DMPs in non-activated preconditioned vs non-preconditioned (*ATP-hyper* and *ATP-hypo*) and activated preconditioned vs non-preconditioned (*Conditioning-hypo* and *Conditioning-hyper*) in ChromHMM categories of monocytes (Roadmap Epigenomics Project). Fisher's exact tests were calculated using all the positions annotated in the EPIC array as background.
